# Comparative analysis of Pearson and Canonical correlation-based functional connectivity matrices for neuroimaging classification tasks

**DOI:** 10.1101/2024.04.23.590747

**Authors:** Ekaterina Antipushina, Maria Zubrikhina, Ruslan Kalimullin, Nikolay Kotoyants, Maxim Sharaev

## Abstract

Machine learning (ML) methodologies offer significant potential for addressing the intricate challenges inherent in the analysis of neuroimaging data within the realm of neurological research. Nonetheless, the effective application of these techniques is markedly contingent upon the particular task and dataset under examination, and the absence of standardized methodologies poses impediments to cross-study result comparisons. This study contributes substantively to the collective endeavor by conducting a comprehensive evaluation and comparative analysis of ML models in the context of predicting schizophrenia and autism spectrum disorder (ASD) utilizing distinct functional Magnetic Resonance Imaging (fMRI) datasets. In this research, we introduce Canonical Correlation Analysis (CCA) as an innovative modality to augment the classification of these multifaceted neurological conditions. By elucidating the efficacy of CCA in ameliorating classification accuracy within the framework of Support Vector Machines (SVM), our study endeavors to propel the domain of neuroimaging and deepen our understanding of these intricate neurological disorders.

## I. Introduction

Functional Magnetic Resonance Imaging (fMRI) has emerged as a pivotal tool within the realm of neuroscience research, affording non-invasive elucidation of cerebral activity and functional interconnections. The proliferation of extensive fMRI datasets has instigated the pursuit of more intricate analytical techniques for the extraction of meaningful insights from this voluminous neuroimaging data, encompassing both the tridimensional spatial dimensions (x, y, z) and the temporal dimension (t). Recent years have witnessed the convergence of machine learning and deep learning paradigms into the realm of fMRI data analysis, showcasing considerable potential in unraveling profound cognizance of cerebral function and temporal cognitive processes.

Machine learning techniques encompass a diverse spectrum of algorithms amenable to fMRI data analysis. These methodologies particularly excel in classification tasks [1], [2], where they discern latent patterns and interrelationships within the data, which may elude conventional statistical approaches. Notably, the application of support vector machines (SVMs) and random forests has found utility in the classification of neurological or psychiatric disorders, effectively distinguishing between healthy subjects and those afflicted with such conditions based on their patterns of brain activation.

Our primary focus in this extensive investigation centers on identifying the most effective approach for predicting neurological disorders, specifically schizophrenia and autism, using a distinct dataset of fMRI data. Our research is anchored in a comparative analysis that establishes a benchmark for traditional machine learning methods. Through the evaluation of their performance, our objective is to shed light on the optimal strategy for fMRI data analysis, thereby driving progress in the exploration of cerebral function and the comprehension of neurological disorders.

Within conventional fMRI data analysis, the foundational step often involves extracting Region of Interest (ROI) information from the original fMRI data. However, the conventional approach of constructing ROI correlation matrices using Pearson correlation coefficients may not be ideally suited for capturing complex relationships between brain regions and can not detect sophisticated patterns of functional connectivity that may be missed by Pearson orrelation [3].

In our research, we introduce a preprocessing method that employs Canonical Correlation Analysis (CCA) for fMRI data, addressing the limitations of traditional preprocessing approaches [4]. We illustrate how our approach enhances the capacity to scrutinize functional connectivity within the cerebral domain, consequently elevating the performance of Support Vector Machine (SVM), SVM in conjunction with Principal Component Analysis (PCA). This enhancement results in improved accuracy in disease classification, marking a notable advancement in our investigation.

## II. Materials and methods

### A. Neurological datasets description

This section provides an overview of the two fMRI datasets used in our study, each consisting of distinct sets of connectivity matrices derived from different brain regions or connectivity methods. These matrices represent brain regions of interest (ROIs) along rows and columns, with the connectivity strength between a given pair of ROIs stored in the cell where these two regions intersect [5].

The first dataset is referred to as The Center for Biomedical Research Excellence (COBRE) dataset [6] with the Automated Anatomical Labeling (AAL) atlas [7]. It consists of 152 connectivity matrices, each with dimensions of 116×116, and includes individuals aged between 18 and 65 in each group. Among the participants, there are 79 individuals diagnosed with schizophrenia, comprising 20 females and 59 males, and 73 healthy controls, including 23 females and 50 males. The age distribution in the normal control group is as follows: Mean age = 36 years, with a standard deviation of ±12 years. In the schizophrenia group, the age distribution is Mean age = 38 years, with a standard deviation of ±14 years. These matrices provide a powerful representation of brain network organization and communication patterns, shedding light on how different brain regions cooperate during specific cognitive tasks or experimental conditions.

The second dataset is the Autism Brain Imaging Data Exchange (ABIDE) [8] with the Craddock200 atlas [9]. It comprises 1035 connectivity matrices, each sized at 200×200 includes 530 individuals diagnosed with ASD (467 males and 63 females) and 505 healthy controls (443 males and 62 females). The age distribution among male subjects with ASD is as follows: Mean age = 17.5 years, with a standard deviation of ±8 years. Among female subjects, the age distribution is Mean age = 16 years, with a standard deviation of ±7 years, and the age range spans from 6.5 to 40 years. The dataset is specifically tailored for the study of ASD and provides an essential collection of connectivity patterns between 200 brain regions, facilitating a comprehensive exploration of the neural correlates associated with this complex neurological condition.where, ROIa - the first region of interest from the corresponding zone, ROIb - second region of interest from the corresponding zone, *t* - time of the experiment, *l1* - components obtained after SVD, accounting for 90% variance.

### B. Functional Connectivity Analysis Using CCA and PCA

In this section, we detail our approach to investigating functional connectivity between distinct brain regions using fMRI data. We employed Canonical Correlation Analysis (CCA) to quantify the strength of interregional relationships and uncover potential connections in fMRI-based functional connectivity analysis. Our study leveraged a comprehensive dataset sourced from COBRE. Our original dataset comprised fMRI scans from a single patient, with each scan containing information pertaining to 116 distinct brain regions, referred to ROI. For each ROI, we possessed a time series of dimensions (150, 116), where 150 represented temporal intervals, and 116 denoted number of ROIs. To address high-dimensional fMRI data challenges and prevent overfitting, we applied Principal Component Analysis (PCA), reducing dimensionality and mitigating multicollinearity among regional time series. Following PCA, we explored functional connectivity via Canonical Correlation Analysis (CCA) with l1 number of components, identifying robust linear relationships that unveiled vital interregional brain connections.

For further analysis and utilization in machine learning, we aggregated the correlation matrices obtained for each patient. Subsequently, we conducted additional data preprocessing, including standardization and normalization, to prepare the data for training and evaluation of ML models (see Figure 1).

**Fig. 1:**
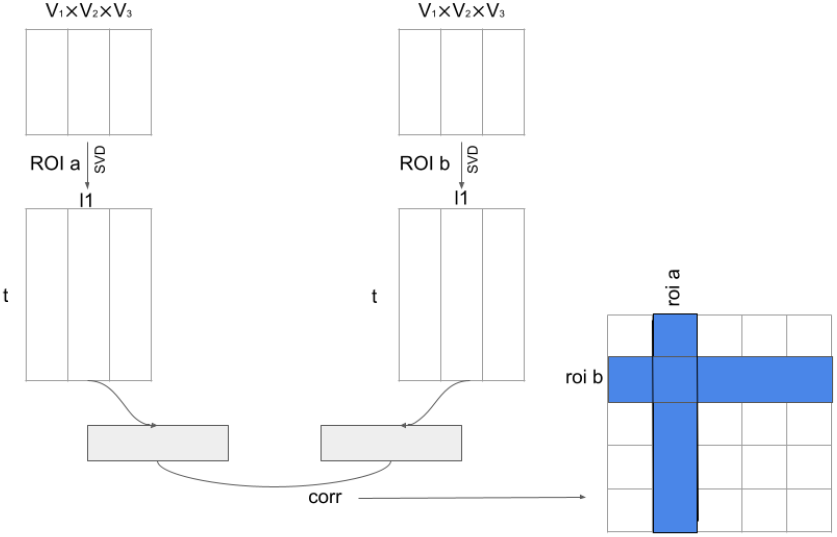
Preprocessing pipeline with Canonical Correlation Analysis where, ROIa the first region of interest from the corresponding zone, ROIb - second region of interest from the corresponding zone, *t* - time of the experiment, *l1* - components obtained after SVD, accounting for 90% variance.

### C. Machine Learning models description

In our research, we utilized various machine learning models to effectively analyze and extract meaningful insights from the connectivity matrices. The selection of these models was driven by their ability to handle the unique characteristics of the data, including its symmetric nature (where the correlation between ROI A and ROI B is the same as between ROI B and ROI A) and multi-dimensional structure.

We considered four different machine learning models: Logistic Regression (LR), Random Forest (RF), Support Vector Machine (SVM), and XGBoost. Logistic Regression was chosen for its simplicity and interpretability, making it well-suited for binary classification tasks. Random Forest was selected due to its capability to handle high-dimensional data and resistance to overfitting. Support Vector Machine was considered for its ability to handle non-linear relationships in the data and its strong generalization properties. Finally, XGBoost was included as a powerful ensemble model known for its excellent performance on structured data and ability to capture complex interactions within the connectivity matrices. To optimize the performance of each model, we conducted a grid search approach with 5-fold cross-validation on a concatenated dataset comprising all connectivity matrices and their corresponding targets, where the targets represented patients with the disorder and healthy individuals. During this process, we fine-tuned the hyperparameters of each model, such as the regularization strength *C* for LR and SVM, the number of estimators n_estimators for RF and XGB, and the learning rate *η* for XGB. This allowed us to identify the optimal configuration for each model, ensuring robust and accurate results in our analysis.

Our primary goal was to determine the best-performing model based on the mean F1 score across the validation data folds. This comprehensive evaluation provided valuable insights into the strengths and limitations of each model in capturing brain network dynamics and identifying neurobiological patterns from fMRI data. After selecting the best model based on the validation data, we further evaluated its performance on independent test splits to assess its generalization capability (see Figure 2).

**Fig. 2:**
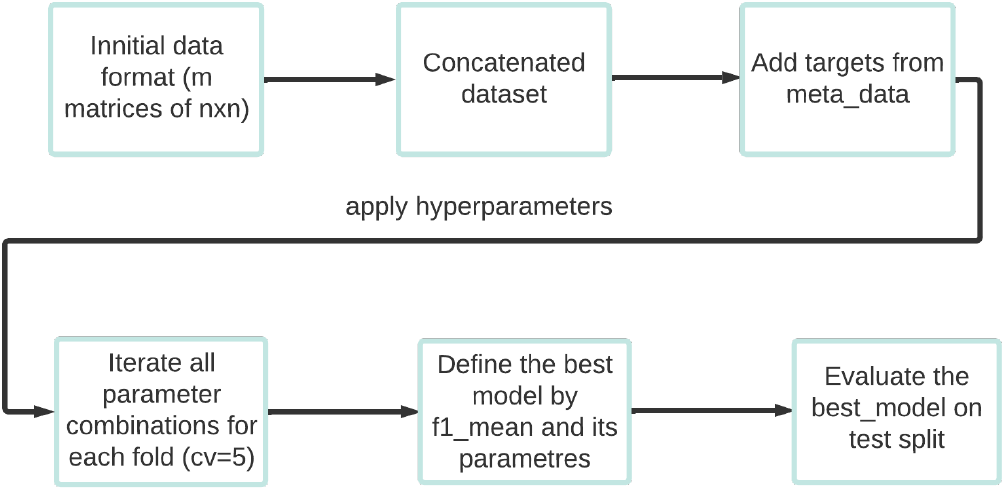
Schematic Representation of ML Model Pipeline

## III. Results

In this research, an investigation was conducted on the COBRE and ABIDE datasets, widely used in functional magnetic resonance imaging (fMRI) analysis. The goal was to evaluate the performance of various classification methods for fMRI data using an 80-10-10 data split (train-validation-test).

The results were presented in Table 1 for the COBRE dataset and Table 2 for the ABIDE dataset. Additionally, Figures 3 and 4 displayed the test scores obtained by deep learning models on the COBRE and ABIDE datasets, respectively.

**TABLE I:** Scores obtained for test splits of COBRE dataset.

| Models | F1 Score | Accuracy | ROC-AUC |
| --- | --- | --- | --- |
| Random Forest | 0.65±0.073 | 0.69±0.064 | 0.69±0.065 |
| XGBoost | 0.67±0.044 | 0.72±0.029 | 0.71±0.033 |
| SVM + PCA | 0.71±0.099 | 0.76±0.062 | 0.75±0.072 |
| Logistic Regression | 0.73±0.056 | 0.76±0.045 | 0.76±0.049 |
| **SVM** | **0.76±0.058** | **0.78±0.039** | **0.79±0.047** |

**TABLE II:** Scores obtained for test splits of ABIDE dataset.

| Models | F1 Score | Accuracy | ROC-AUC |
| --- | --- | --- | --- |
| Random Forest | 0.65±0.037 | 0.67±0.030 | 0.67±0.031 |
| XGBoost | 0.65±0.007 | 0.63±0.010 | 0.63±0.011 |
| Logistic Regression | 0.66±0.013 | 0.66±0.013 | 0.66±0.013 |
| SVM | 0.67±0.012 | 0.67±0.013 | 0.67±0.013 |
| **SVM + PCA** | **0.69±0.017** | **0.68±0.015** | **0.68±0.016** |

Remarkably, the Support Vector Classifier (SVC) consistently achieved the highest scores on both datasets. This contributes to the existing knowledge in the field and can assist researchers and practitioners in selecting appropriate classification methods for fMRI data analysis. The study emphasizes the importance of data partitioning strategies when assessing the performance of classification models on fMRI datasets.

Through a comparison with state-of-the-art approaches presented by Kamalaker Dadi et al. [10] and Anibal Sólon Heinsfeld et al. [11], we demonstrated the effectiveness of classic machine learning models, including the Support Vector Classifier (SVC), in handling fMRI data classification tasks. Despite the relatively small size of the datasets, our findings indicate that these classic models can yield competitive results comparable to more complex deep learning approaches. This benchmarking study offers valuable insights into the potential of different models for fMRI analysis and establishes a reference point for future advancements in the field of functional connectome-based predictive models using fMRI data.

Next, we obtained results based on CCA preprocessing. For ML approaches, the best results were achieved using the SVM method, demonstrating a significant increase in data classification accuracy which are presented in Table 3. Our results also outperformed those obtained for Pearson based preprocessing and surpassed the results of state-of-the-art approaches for ROI analysis. This confirmed our hypothesis of CCA as a novel approach to data preprocessing.

**TABLE III:** Scores obtained for test splits of COBRE dataset. CCA matrices.

| Models | F1 Score | Accuracy | ROC-AUC |
| --- | --- | --- | --- |
| XGBoost | 0.61±0.114 | 0.67±0.075 | 0.66±0.084 |
| Random Forest | 0.62±0.148 | 0.73±0.087 | 0.71±0.096 |
| Logistic Regression | 0.68±0.043 | 0.75±0.030 | 0.731±0.033 |
| SVM + PCA | 0.74±0.047 | 0.78±0.030 | 0.78±0.036 |
| **SVM** | **0.77±0.087** | **0.82±0.061** | **0.81±0.068** |

## IV. Discussion

Within this context, we present the core findings arising from our study, with a specific focus on unraveling the determinants that account for the superior scores reached by ML models, such as Support Vector Machine with Principal Component Analysis (SVM+PCA) on Pearson’s functional connectivity matrices. Additionally, we explore the relationship between sample size and model performance quality and investigate the impact of feature selection on model performance, particularly when dealing with small datasets.

In addition to the previously mentioned methods, we estimated scores on functional matrices with CCA preprocessing step. This approach yielded favorable results when combined with SVM, SVM+PCA resulting in improved prediction accuracy. However, it’s important to note that CCA did not demonstrate any accuracy improvements when applied in conjunction with other classifiers. This discrepancy could be attributed to the fact that CCA introduces additional features to our dataset without a corresponding increase in data volume, potentially leading to a decrease in accuracy.

In our work, we took a comprehensive approach by utilizing all region-of-interests (ROIs) for fMRI analysis. This allowed us to capture a broad representation of brain activity and connectivity patterns. However, it is worth exploring the impact of focusing only on important features, especially when investigating specific conditions like Autism Spectrum Disorder (ASD) or Schizophrenia. Not all ROIs may contribute equally to the classification task, and some ROIs could be more informative or relevant for distinguishing between healthy individuals and those with the respective conditions.

Conducting a feature importance analysis or employing feature selection techniques could help identify the most discriminative ROIs for ASD/Schizophrenia classification. By focusing on the most relevant features, you may enhance the performance of your classification models, improve interpretability, and potentially uncover specific brain regions or functional connections associated with the conditions.

## V. Conclusions

In summary, our findings underscore the significance of model simplicity and judicious feature selection when dealing with limited datasets in the context of neuroimaging studies. Additionally, the outcomes derived from the application of Canonical Correlation Analysis (CCA) to our data and models yield valuable insights. We conducted a benchmark that involved a thorough comparison between models employing CCA and those using conventional data analysis methods. This rigorous evaluation allowed us to elucidate situations in which CCA exhibited superior performance relative to standard methodologies.

